# Genome scale CRISPR Cas9a knockout screen reveals genes that controls glioblastoma susceptibility to the alkylating agent temozolomide

**DOI:** 10.1101/2021.08.11.455972

**Authors:** Chidiebere U Awah, Jan Winter, Olorunseun O. Ogunwobi

**Affiliations:** Department of Biological Sciences, Hunter College of The City University of New York, New York, NY, USA; Joan and Sanford I. Weill Department of Medicine, Weill Cornell Medicine, Cornell University, New York, NY, USA; Hunter College for Cancer Health Disparities Research, Hunter College of The City University of New York, New York; Steinbrenner Laborsysteme GmbH, Wiesenbach, Germany

**Author notes:** Correspondence should be addressed to, Chidiebere U Awah MD MSc PhD, Department of Biological Sciences, Hunter College of The City University of New York, New York, NY, USA.

## Abstract

Glioblastoma is the most fatal of all primary human brain tumors with 14 months survival, at best. The mainstay therapy for this tumor involves temozolomide, surgery, radiotherapy and tumor treating electric field. Cancer resistance to commonly available chemotherapeutics remains a major challenge in glioblastoma patients receiving treatment and unfavorably impact their overall survival and outcome. However, the lack of progress in this area could be attributed to lack of tools to probe unbiasedly at the genome wide level the coding and non-coding elements contribution on a large scale for factors that control resistance to chemotherapeutics. Understanding the mechanisms of resistance to chemotherapeutics will enable precision medicine in the treatment of cancer patients.

CRISPR Cas9a has emerged as a functional genomics tool to study at genome level the factors that control cancer resistance to drugs. Recently, we used genome wide CRISPR-Cas9a screen to identify genes responsible for glioblastoma susceptibility to etoposide. We extended our inquiry to understand genes that control glioblastoma response to temozolomide by using genome scale CRISPR. This study shows that the unbiased genome-wide loss of function approach can be applied to discover genes that influence tumor resistance to chemotherapeutics and contribute to chemoresistance in glioblastoma.

## Introduction

Glioblastoma is the deadliest of all primary brain tumors with a very poor survival outcome. Temozolomide, an alkylating agent, is used as a mainstay drug in treating of glioblastoma patients along with radiation, tumor treating field and surgery^1,2^. However, this drug offers little to no benefit for glioblastoma patients. Hence, the objective of the generation of this data is to understand and uncover genes that control glioblastoma susceptibility and resistance to temozolomide using an unbiased genome scale CRISPR Cas9a knockout screen.

## Material and Methods

### Details of Source of all samples

#### Cell lines

The glioma cell line, U251 was commercially obtained from ATCC and used in the genome scale CRISPR screen.

#### Reagent

The U251 cells were grown in 10% FBS DMEM media with antibiotics up to 80% confluency before downstream analysis.

#### CRISPR Knockout library

The Brunello library pooled gRNA (Cat no: 73178-LV) were commercially purchased from Addgene. This library contains 70,000sgRNA, with 60,000 guides targeting approximately the 20,000 genes in the human genome at the rate of 3-4 guides per gene. There are about 10,000 guides which are the non-targeting controls.

#### Determination of viral titre and multiplicity of infection

3×10^6^ of U251 cells are seeded into 12-well plate in 2ml. The Brunello library as virus supernatant were added at 400μl, 200μl, 100μl, 75μl,50μl, 25μl and 8μg/μl of polybrene is and spinfected at 1000g at 33°C for 2hrs. Cells then are incubated at 37°C. After 24hrs the cells are harvested and seeded at 4×10^3^ with puromycin for 96hrs with a well containing cells that were not transduced with any virus. After 96hrs the cell titre glo (Promega cat is used to determine cell viability at MOI 21%. At the multiplicity of infection (MOI) of 21% we are able to infect 1 sgRNA/cell (**Figure 1A-C**).

**Figure 1:**
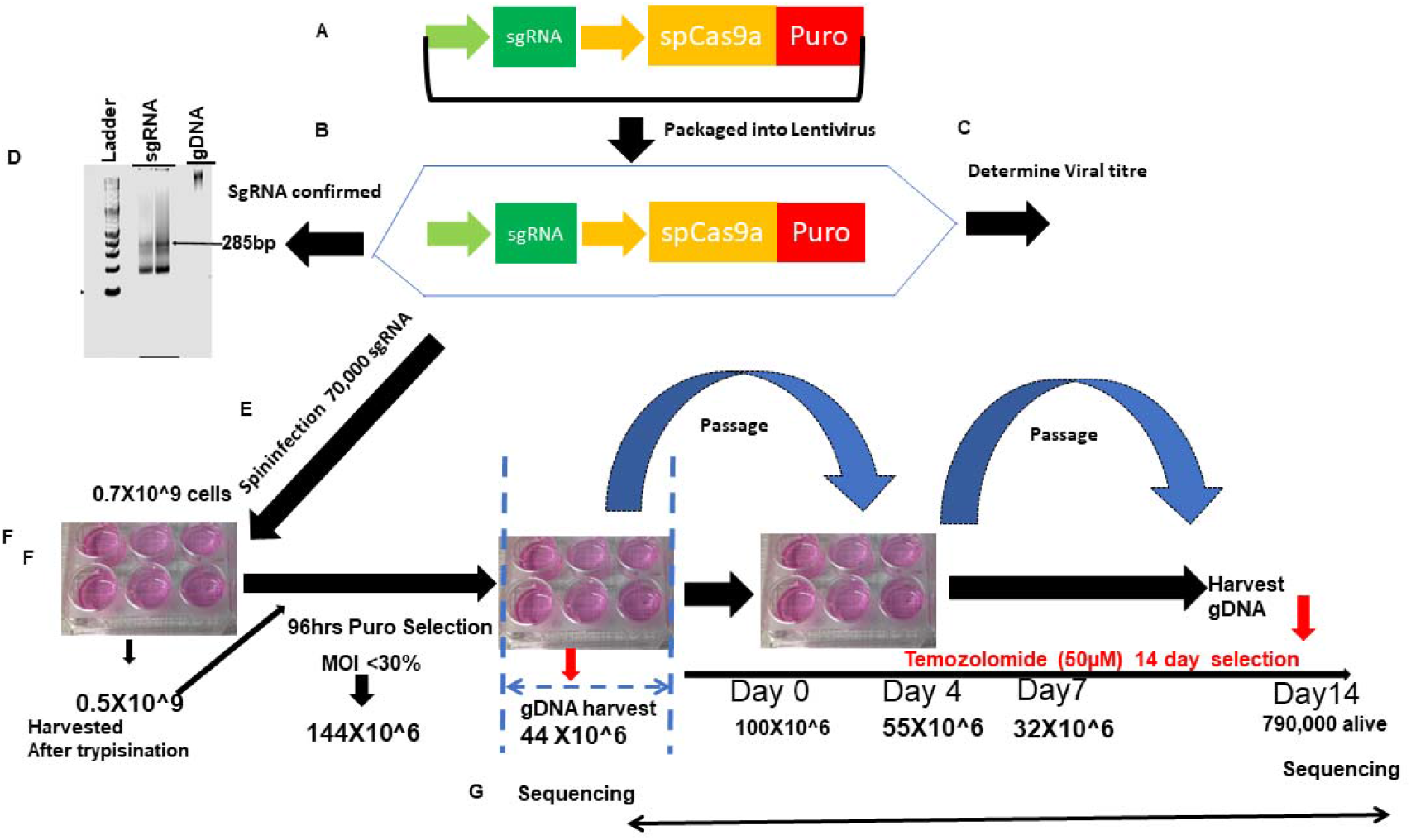
Strategy for genome scale CRISPR screen. A. shows the schematic depiction of the sgRNA library B. Library is packaged into lentiviral. C viral titre and MOI is determined. D A gel shows amplification of sgRNA library. E Lentivirus containing sgRNA is spinfected into glioma cells. F Transduced cells are selected with puromycin after which some cells are harvested for baseline sgRNA representation, the rest of the cells are placed under the treatment of temozolomide/DMSO/puromycin for 14 days. G. cells are harvested, sgRNA extracted and sequenced on NEXT generation sequencers.

#### Genome wide scale CRISPR-Cas9a knockout screen

To do the CRISPR screen, we expanded the U251 to 500million cells and then spinfected with 70,000sgRNA. The library was integrated into the cells by spinfection, and subsequently selected with 0.6μg/ml of puromycin for 4days. This selection is aimed at the cells that have been rightly integrated with the sgRNA that incorporates the puromycin cassette into their genome. We achieved an MOI of 21% in two independent screens. At the end of day 4, about 150million cells survived the selection. We used 50 million of selected cells for the extraction of genomic DNA. The base sgRNA representation is obtained by amplification of the sgRNA with unique barcoded primers. The remaining 100million cells were expanded for 2days once cells grew to 300 million. 100 million of cells were treated with temozolomide at concentration of 50 μM for 14 days, and the 100 million each were treated with DMSO and Puromycin for 14 days respectively and served as control. DMSO is control for temozolomide treated cells, whereas puromycin is for identification of essential genes driving glioma After 14days, the cells were harvested, the gDNA extracted, and the sgRNA amplified with another unique barcoded primer (**Figure 1D-G; Figure 2 A, Figure 3 A-B**).

**Figure 2:**
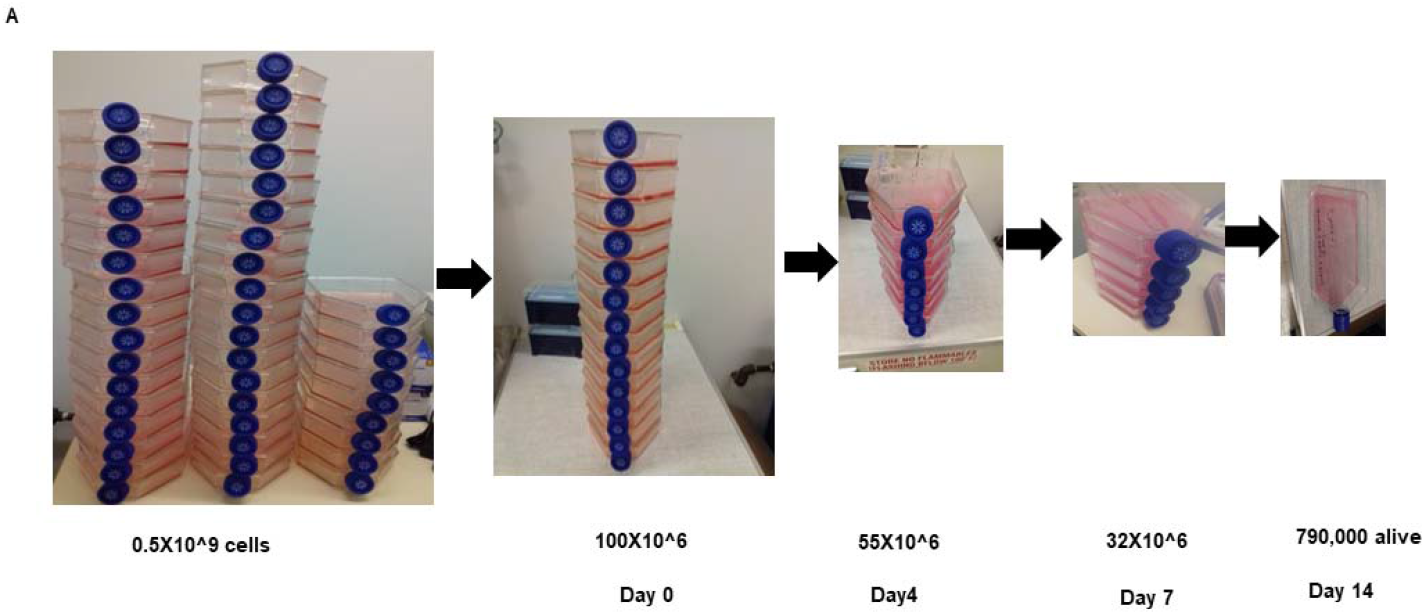
Kinetics of cell selection pressure under chemotherapy in genome scale CRISPR screen. A. shows the schematic depiction of number of T225 flask of cell starting from over half a billion cells to the number that remains after 14 days of selection.

**Figure 3:**
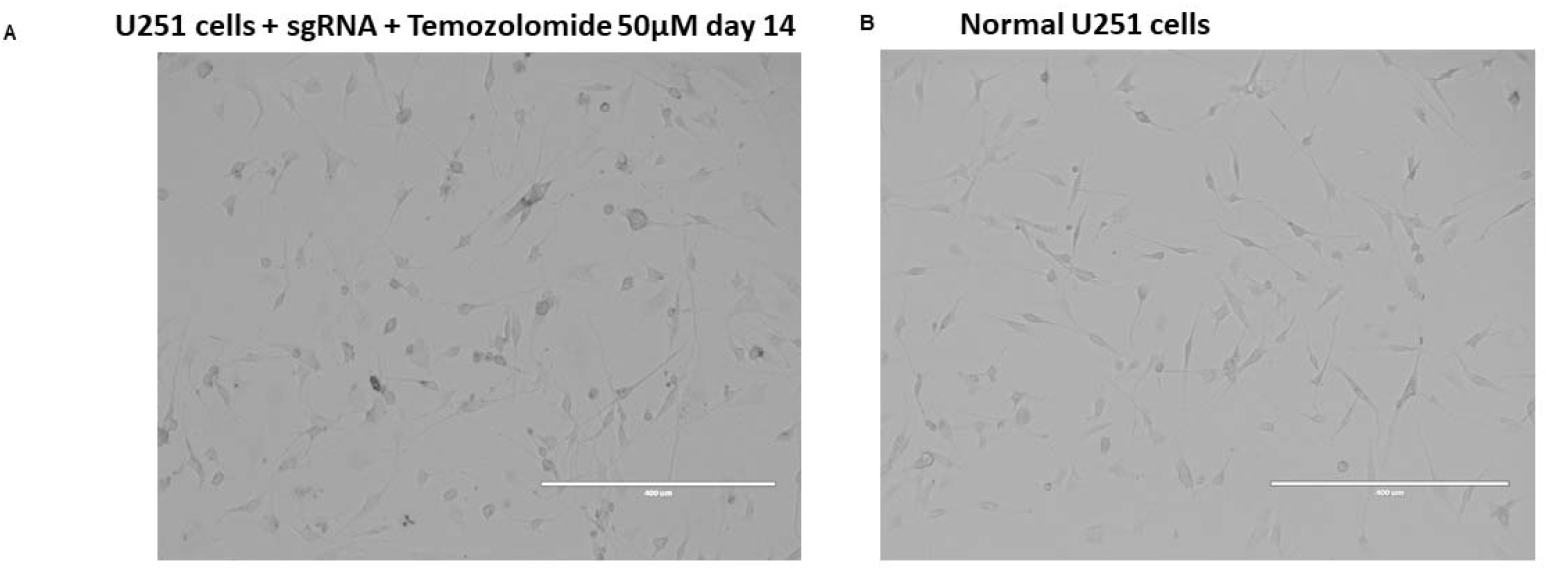
Morphology of cells transduced with CRISPR library under drug selection compared to wildtype control. A. shows U251 cells transduced with sgRNA treated with temozolomide for 14 days and B shows normal U251 wildtype controls.

#### DNA extraction and PCR amplification of pooled sgRNA

The genomic DNA (gDNA) were extracted with the Zymo Research Quick-DNA midiprep plus kit (Cat No: D4075). gDNA was further cleaned by precipitation with 100% ethanol with 1/10 volume 3M sodium acetate, PH 5.2 and 1:40 glycogen co-precipitant (Invitrogen Cat No: AM9515). The gDNA concentration were measured by Nano drop 2000 (Thermo Scientific). The PCR were set up as:

##### 50μl reaction

5μl of 10x buffer
4μl of 50mM DNTP (1/25)
0.5μl of P5 forward primers equimolar mix
10μl of 20μM reverse primer (1 per experimental condition)
gDNA (20ng for sgRNA library) or 2-10μg for sgRNA in various the experimental condition
1.5μl of Extaq enzyme
27μl of H20

##### With NEBNEXT PCR polymerase enzyme the following recipe is used

25μl of NEB Next High Fidelity Master Mix (2x)
1μl of gDNA (20ng for sgRNA library) or 2-10μg for sgRNA in various the experimental condition 1.25μl of P5 forward primers equimolar mix
1.25μl of 20μM reverse primer (1 per experimental condition)
21.5μl ultrapure H20
The sequences of P5 forward primers and the reverse primers can be found described^14^.

##### The PCR cycle used is

**The PCR cycle used is:**
| Cycle number | Denature | Anneal | Extend |
| --- | --- | --- | --- |
| 1 | 98°C 3mins |  |  |
| 2-23 | 98°C 10sec | 63°C 10sec | 72°C 2mins |

Don’t exceed 23 cycles. We have found that difficulty gDNA containing sgRNA can be amplified using annealing temperature gradient of 53-68°C. The sgRNA knockout size is 285bp. All extracted gDNA from the cells were all amplified to maintain the library coverage.

#### Next Generation Sequencing

The sgRNAs amplified and gel extracted from **Figure 4** were pooled together and sequenced in a Next generation sequencer (Next Seq) at 300 million reads for the four sgRNA pool aiming at 1,000reads/sgRNA. The samples were sequenced according to the Illumina user manual with 80 cycles of read 1 (forward) and 8 cycles of index 1^3,4^. 20% PhiX were added on the Next Seq to improve library diversity and aiming for a coverage of >1000reads per SgRNA in the library.

**Figure 4:**
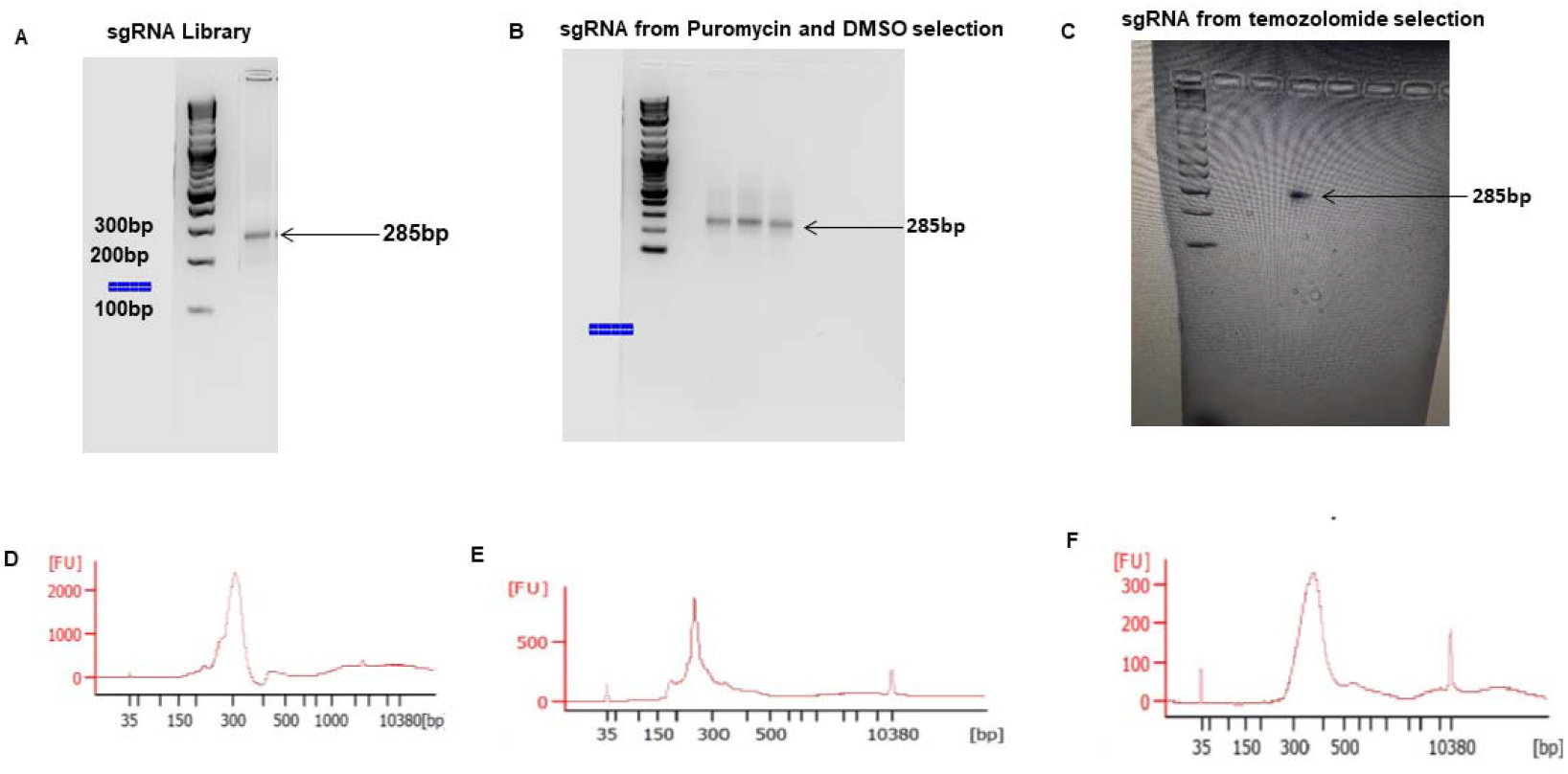
PCR amplification of sgRNA from library, puromycin, DMSO and temozolomide selected glioblastoma: A-C shows agarose gel pic of the sgRNA amplified from library, puromycin, DMSO and temozolomide treated cells. D-F shows electropherogram measurement of the amplified sgRNA from the different conditions.

#### CRISPR screen data analysis

All data analysis was performed with the bioinformatics tool CRISPR Analyzer (http://crispr-analyzer.dkfz.de/)^5^.

The sequence reads obtained from Next Seq as fastq files were aligned with human genome in quality assessment to determine the percentage that map to the total human genome (**Figure 5**).

**Figure 5:**
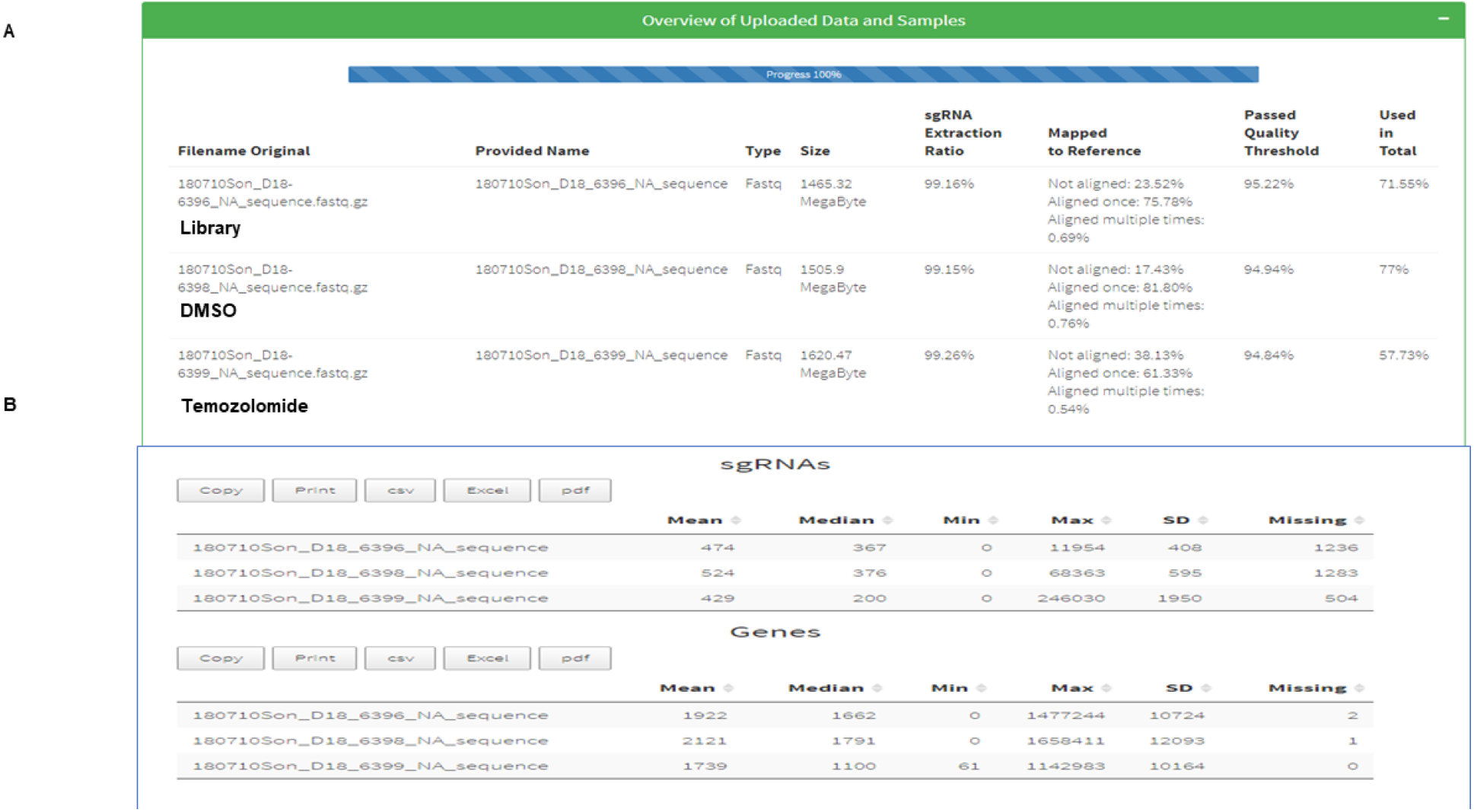
Alignment of Next Seq sgRNA reads from library, DMSO and Temozolomide treated cells. A show the mapping of sgRNA library (75.78%) DMSO (81.8%) and temozolomide (61.33%) mapped to the human reference genome. B. shows the number genes and sgRNA missing from the mapped library which were few.

To find genes that were enriched or depleted under various treatment condition, the CRISPRAnalyzer performs four rigorous statistical analysis where are (1) DeSeq 2 (Log2 fold sgRNA count) 3 (Wilcoxon t-test) and 4 (Z-ratio) comparing the enrichment of guides in treatment conditions versus their non-targeting controls. P value of 0.05 is used as a cut off for guides statistical enrichment. The sgRNA that does not meet a read count of 20 is removed. Hit calling from the CRISPR screen was done based on sgRSEA enriched, p<0.01 was used for significance based on Wilcoxon test.

To set up the analysis, the sgRNA reads (library, puromycin, DMSO and temozolomide) replicates were loaded unto the software. The sgRNA that does not meet a read count of 20 is removed. Hit calling from the CRISPR screen was done based on sgRSEA enriched, p<0.01 was used for significance based on Wilcoxon test.

CRISPR Analyzer does also correlation analysis based on Pearson and Spearman statistical analysis to determine the relatedness between samples replicates and then uses log2 fold SgRNA read count to calculate the cumulative frequency between guides in various treated samples compared to controls in cumulative frequency curve (**Figure 6**) and in scatter plot (**Figure 7**).

**Figure 6:**
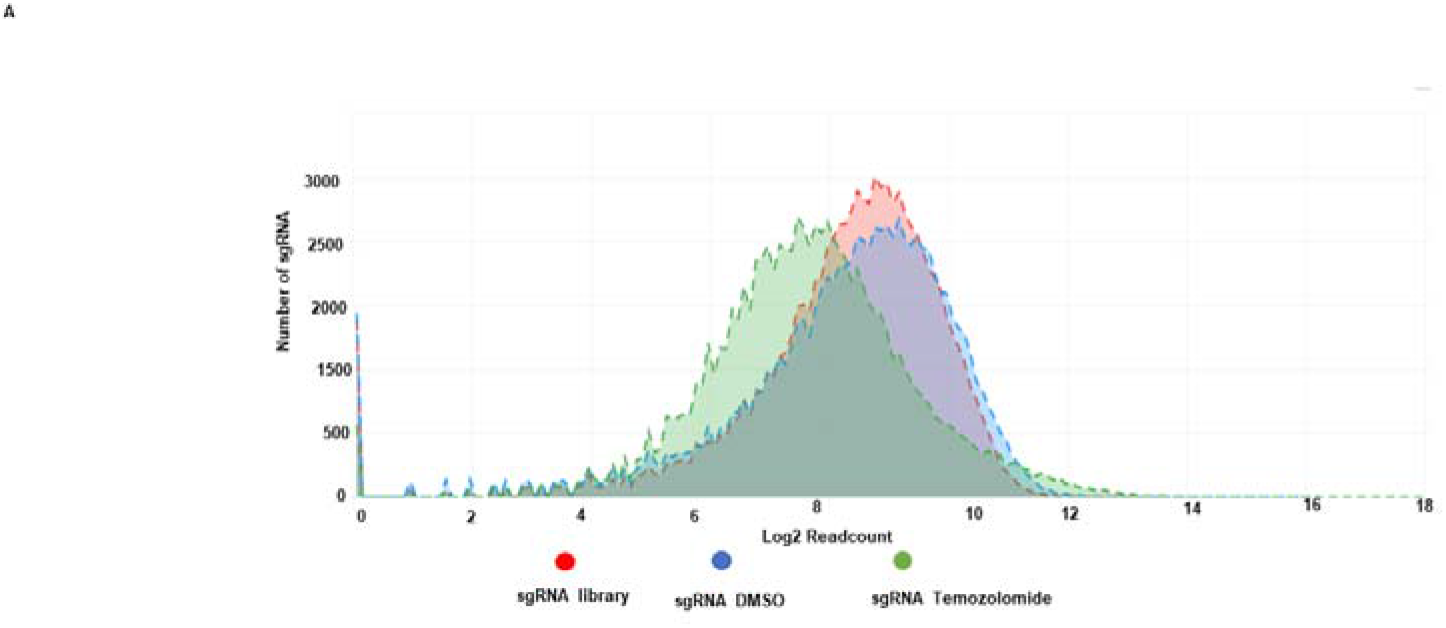
Cummulative frequency of sgRNA distribution in CRISPR library, DMSO and Temozolomide treated glioblastoma cells. A shows cumulative frequency distribution of log 2 sgRNA read count in sgRNA library (red), sgRNA DMSO (blue) and sgRNA Temozolomide (green).

**Figure 7:**
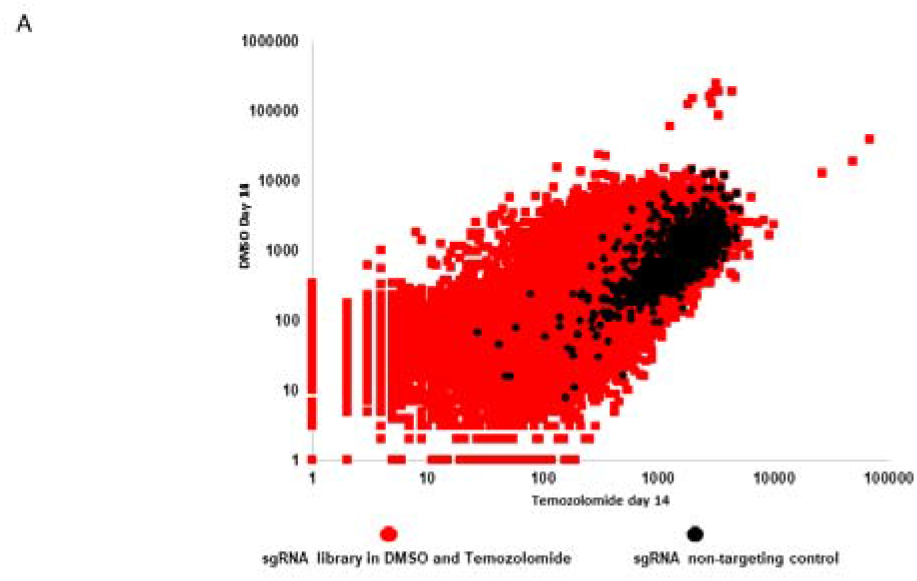
Scatter plot distribution of sgRNA in DMSO versus temozolomide treated glioblastoma cells. **A** Scatter plot depicting sgRNA distribution in DMSO (y-axis) and Temozolomide (x-axis) and their non-targeting controls (black dots).

### Dataset validation

To validate the data from the screen, we found that NSUN6 a 5mC RNA methyltransferase is enriched in the CRISPR screen under temozolomide compared to DMSO and puromycin selection for 14 days. We performed a single gene editing of NSUN6 and proved that loss of NSUN6 led to loss of 5mC and caused resistance to temozolomide. This data has been presented in the primary article ^6^.

## Data availability

All data underlying the results are available as part of the article and no additional source data are required^7^.

**DOI**: 10.5281/zenodo.5167143

Data 1: sgRNA library

Data 3: DMSO treated sgRNA library replicate 1

Data 4: DMSO treated sgRNA library replicate 2

Data 5: Puromycin treated sgRNA Library replicate 1

Data 6: Puromycin treated sgRNA library replicate 2

Data 7: Temozolomide treated sgRNA library

## Software availability

CRISPRAnalyzer the software used in this data analysis is publicly available at (http://crispr-analyzer.dkfz.de/).

## Acknowledgment

We acknowledge our colleagues who provided support in the cell culture.

